# Rapid and visual detection of *Schistosoma japonicum* circulating nucleic acids by CRISPR-Cas12a

**DOI:** 10.1101/2023.07.31.551221

**Authors:** Qian Li, Shuqi Yang, Miao Cheng, Chen Xing, Xiaofeng Wang, Yating Zhu, Yunshu Tang, Wenhui Cheng, Jijia Shen, Yinan Du

## Abstract

**Background:** Schistosomiasis is the second most destructive parasitic disease globally. It poses a serious threat to human health if not diagnosed and treated promptly. However, the current detection methods have limitations due to their dependence on specialized equipments and personnel, and the long detection time, resulting in less-than-ideal detection outcomes. Therefore, more accurate and sensitive assays are required to aid in the control of schistosomiasis. This study aims to develop a rapid and practical method for detecting *Schistosoma japonicum* circulating nucleic acid by combining the Cas12a protein in the CRISPR (Clustered regularly interspaced short palindromic repeats) system with recombinase-mediated isothermal nucleic acid amplification technology.

**Methodology/Principal Findings:** We created a CRISPR-Cas12a detection method for Schistosomiasis. *SjR2* and *SjCHGCS19* genes with repetitive copies *in the Schistosoma japonicum* genome were targeted for detection. The CRISPR-Cas12a detection method can achieve a detection sensitivity of 1 cp/µL, with no cross-reaction with the other three human-infecting trematode parasites. To vividly simulate clinical patients at different infection stages, we collected rabbits’ serums at 1 d, 3 d, 1 w, 3 w, 5 w, and 7 w post-infection. Circulating *Schistosoma japonicum* DNA in the serum can be detected as early as 3 days post-infection. The sample and positive rates of each stage were analyzed and the CRISPR-Cas12a detection method prevailed to Nested-PCR. Additionally, our assay offers flexible results readout options, including lateral test strips.

**Conclusions/Significance:** This CRISPR-Cas12a detection method enables sensitive and specific detection of Schistosomiasis by detecting highly repetitive *SjR2* and *SjCHGCS19* genes. The method was fully validated with rabbits’ serums collected at different infection stages, which can be applied to monitoring the early infection of schistosomiasis in resource-limited regions.

**Author Summary:** With the development of schistosomiasis prevention and control efforts in recent years, the prevalence rate and infection intensity of schistosomiasis have decreased, making early and accurate diagnoses more challenging. Combining CRISPR-Cas12a protein with isothermal amplification technology, a detection method for the multi-copy sequences *SjR2* and *SjCHGCS19* in the genome of *Schistosoma japonicum* was developed in this study. The results demonstrated that the detection sensitivity of this method could reach 1 cp/µL, with no cross reactivity with three other fluke-like human parasites. In this study, serum samples from 15 rabbits infected with *Schistosoma japonicum* were used to validate the method. The results demonstrated that rabbits could be identified as early as the third day after infection. CRISPR-Cas12a detection method has a higher positive rate than traditional Nested-PCR. In addition, lateral test strips enable direct observation of our assay results. The aforementioned research findings indicate that the CRISPR-Cas12a-based detection method can be utilized for the early diagnosis of schistosomiasis. In conjunction with the result reading method of the lateral test strips, it also provides a new method for the on-site diagnosis of schistosomiasis in environments with limited access to resources.

## Introduction

Schistosomiasis is a zoonotic parasitic disease induced by infection with schistosomes[1]. Schistosomiasis is prevalent in 78 countries and regions worldwide, infecting approximately 250 million individuals[2, 3]. It is a fatal parasitic disease that is frequently ignored[4]. Schistosomiasis, if not diagnosed and treated in a timely manner, will negatively impact the health, growth, and development of children, as well as place a significant economic burden on the families of afflicted individuals and have a negative impact on economic and social developments[5, 6]. Researchers have developed a variety of schistosomiasis diagnostic methods. The pathogenic diagnostic method involves the observation of schistosome eggs in the stool, urine, and intestinal biopsy samples through a microscope[7]. However, this method has limitations, such as unstable egg excretion in patients with a mild or early-stage infection that may lead to misdiagnoses [8]. The immunological diagnosis, which relies on the antigen-antibody reaction to detect the disease, has high sensitivity but suffers from poor specificity due to cross-reactivity with other worms[9–11]. Molecular biology diagnostic methods, such as polymerase chain reaction (PCR), can detect DNA of schistosome in blood samples infected with schistosomes and have high sensitivity and specificity[12]. However, the dependence on sophisticated instruments and trained operators limits their extensively implemented in the field. Based on the aforementioned research background, this study aims to establish an efficient and specific detection method for the diagnosis of schistosomiasis that is readily applicable in the field.

The Cas12a protein of the CRISPR-Cas system, which can emit non-specific single-stranded DNA trans-cleavage activity when it specifically targets the target double-stranded DNA, has demonstrated the tremendous potential for the implementation in the field of molecular diagnosis[13, 14]. For the rapid and specific nucleic acid detection, CRISPR-Cas12a is typically combined with Recombinase Polymerase Amplification (RPA) [15], commonly referred to as DNA endonuclease-targeted CRISPR trans reporter (DETECTR) [16]. The target DNA is rapidly amplified by RPA at a constant temperature of 37 ℃, providing sufficient amounts of target DNA to activate Cas12a to cleave fluorescent single-stranded DNA reporter. The cleaved reporter will free fluorophore moiety and report the detection results. The CRISPR-Cas12a detection method has been successfully applied to the detection of bacteria[17, 18], exosomes[19], and parasites[20, 21], among other applications.

In this study, we combined Cas12a protein with Recombinase Aided Amplification (RAA) [22] technology to develop a detection method for *Schistosoma japonicum SjR2* (GenBank accession number AF412221) and *SjCHGCS19* (GenBank accession number FN356221) genes[23, 24]. The *SjR2* and *SjCHGCS19* genes are highly repetitive DNA sequences located on *Schistosoma japonicum* chromosomes that are easily detectable. Also, the detection system was optimized for buffer and probe concentration to increase the detection sensitivity to 1 cp/µL. This study effectively implemented the CRISPR-Cas12a detection method to blood samples from rabbits infected with *Schistosoma japonicum* for the first time, achieving more sensitive detection results than traditional methods. To address the problem of apparatus limitations for on-site detection of schistosomiasis, this study’s test results can be accessed directly via two practical methods. First, the fluorescence of the reaction tube is read with the naked eye under UV; Second, the results are read using a lateral test strip. Through these two flexible readouts, we can provide more accurate on-site detection of schistosomiasis in environments with limited resources and no access to specialized apparatus.

## Materials and methods

### Materials

Primers for Nested-PCR and RAA were synthesized by Tsingke Biotechnology (Nanjing, China). Reporter probe was synthesized by Tsingke Biotechnology (Nanjing, China). LbaCas12a was purchased from New England Biolabs (M0653T). HiScribeT7 Quick High Yield RNA synthesis kit (NEB E2050S) and Monarch RNA Cleanup Kit (NEB T2040L) for in vitro transcription and purification were purchased from New England Biolabs. PrimeStar HS DNA polymerase was purchased from Takara (Dalian, China). The RAA kit was purchased from Hangzhou ZhongCe BioSci&Tech Co. Ltd (Hangzhou, China), and the DNA extraction kit was purchased from TIANGEN (Beijing, China).

## Methods

### Extraction of Schistosoma japonicum genome

The following is the primary stages of phenol-chloroform DNA extraction: First, prepare 25 : 24 : 1 Tris-saturated phenol, chloroform, and isoamyl alcohol. Then, place adult *Schistosoma japonicum* in a 1.5 mL centrifuge tube, add a sufficient amount of DNA lysate (10 µL proteinase K), cover the centrifuge tube with parafilm, and place it in a water bath at a constant temperature (56 ℃, 5 hours). Then, add an equal volume of Tris saturated phenol (500 µL), agitate vigorously, and stir for 10 minutes. Centrifuge (4 ℃, 12000 R, 7 min), and transfer the liquid in the upper layer to a new centrifuge tube. Shake for 10 minutes while adding 450 µL of a mixture of Tris saturated phenol, chloroform, and isoamyl alcohol to the centrifuge tube containing the precipitate. Centrifuge (4 ℃, 12000 R, 7 min). The supernatant is pipetted into a new centrifuge tube, and 400 µL of a mixture of chloroform and isoamyl alcohol (chloroform: isoamyl alcohol = 24 : 1) is added. Centrifuge: (4 ℃, 12000 R, 7 min) Draw the supernatant into a new centrifuge tube and add 2.5 times the amount of absolute ethanol that has been chilled at −20 ℃. The sample was then centrifuged at 12000 R for 7 minutes at 4 ℃. Discard the supernatant, keep the white precipitate (DNA), add 400 µL of frozen 75% ethanol at −20 ℃, and pipette repeatedly to dissolve. Repeat the previous step twice (rinse three times with ethanol at 75% concentration). The extraction of DNA is complete. Maintain at −80 ℃.

### Animal modeling and sample collection

The Animal Centre at Anhui Medical University purchased rabbits for this project’s animal modeling. The *Schistosoma japonicum* used in this study was extracted from the hepatic portal vein of infected rabbits that had been sacrificed. *Paragonimus westermani, Fasciolopsis buski,* and *Clonorchis sinensis* were acquired from Henan Yongzong Science and Education Equipment Co., Ltd. The Experimental Animal Ethics Committee of Anhui Medical University (LLSC20221092, Anhui Medical University) has approved this study.

The schistosomiasis-infected snails originated from the Jiangxi Provincial Institute for Schistosomiasis control. The snails were placed in a container containing water and then placed under a suitable light source and temperature to allow the schistosome cercariae to escape. Twenty rabbits were divided at random into two groups. In the first group, 15 rabbits were infected with 30 cercariae of *Schistosoma japonicum* via the abdominal skin (observed and enumerated using a microscope and a sliding glass), and in the second group, 5 rabbits were not infected to serve as a control. After infection, blood samples were taken from the ear veins of each rabbit on the 1 d, 3 d, 1 w, 3 w, 5 w, and 7 w. After each blood sample was collected, it was deposited in an incubator at 37 ℃ for 1 hour, and then the serum was centrifuged (for 10 minutes at 2000 rpm). All serum samples were frozen at −80 ℃ until use. In this study, after the seventh week of blood collection, the rabbits were sacrificed and dissected, the liver tissue was extracted, and adult *Schistosoma japonicum* was extracted from the portal vein to confirm the actual infection.

### Serum circulating DNA extraction

Cell-Free Circulating DNA (cfDNA) in the blood is a DNA fragment derived from *Schistosoma japonicum*’s partial degradation following the infection of rabbits with *Schistosoma japonicum*. This DNA fragment is free in rabbit blood and can be extracted for the nucleic acid detection[25]. In this study, after comparing several procedures, the magnetic bead method serum/plasma cell-free DNA extraction kit (TIANGEN: DP709) was selected, and the cell-free DNA was extracted according to the instructions.

### Design of Nested-PCR and RAA primers and preparation of crRNA

According to the *SjR2* and *SjCHGCS19* gene sequences in the NCBI database (<u>National Center</u> <u>for Biotechnology Information</u> (nih.gov)), Nested-PCR and RAA primers were designed, and sequence-specific comparisons were undertaken in the database (S1 and S2 Table). The crRNA corresponding to the conserved sequence matching Cas12a (5’-TTTN) was designed after sequence alignment (S3 Table). In order to detect crRNA, it is necessary to first synthesize the single-stranded DNA template of crRNA, then perform in vitro transcription of RNA according to the kit’s instructions, and lastly purify the resulting crRNA. Then, various crRNAs were screened to isolate the crRNA with the highest sensitivity. Using the SnapGene software, RAA primers were designed based on the screened optimal crRNA. Using BLAST and sequence alignment, the specificity of the designed RAA primers and candidate target sequences of the *Schistosoma japonicum* genome was analyzed.

### Construction of recombinant plasmid standards

Nest PCR amplified high-copy sequences in the *Schistosoma japonicum* genome, including gene fragments of *SjR2* and *SjCHGCS19*.Using the TA cloning technique, the gene fragment was fused to the T-Vector pMD19 vector, and the extracted plasmid was transformed.

### Recombinase aided amplification (RAA)

The RAA reaction system was configured in accordance with the kit’s instructions (ZhongCe BioSci&Tech Co., Ltd., Hangzhou, China). 41.5 µL buffer A, 2 µL (10 µM) each for forward primer and reverse primer, 2 µL template DNA, and 2.5 µL buffer B (Magnesium acetate) were added to the reaction system, followed by capping the tube, briefly centrifuging the reaction system, and incubating it at 39 ℃ for 30 minutes.

### CRISPR-Cas12a method for detection of *Schistosoma japonicum* DNA

CRISPR detection is performed step by step. First, *Schistosoma japonicum* DNA was pre-amplified using RAA to generate more target sequence substrates for CRISPR-Cas12a. Next, use CRISPR-Cas12a detection method, detection system configuration: 16 µL of RNase-free water, 2 µL of 10 × NEB buffer r2.1, 1 µL of LbaCas12a (1 µM), 1 µL of crRNA (1 µM) and incubate at 37℃ for 10 minutes, the Cas12a-crRNA complex is formed. Then add 0.1 µL DNA reporter probe (160 µM) to the reaction and mix. Add 2 µL RAA product to the above reaction system and incubate for 40 cycles at 37 ℃ on the Bio-Rad CFX96 Touch real-time PCR detection system. Fluorescence data were collected at 2 min intervals.

### Lateral test strip

The amplified test sample was added to the CRISPR-Cas12a reaction system and incubated for one hour at 37 ℃. After incubation, add 100 µL of 1 × PBS for dilution to the system, and pipette, and add 60 µL to the sample area of the lateral test strips. Observe the results 10 to 20 minutes later.

This study employs the test strip as the line elimination method: both the quality control line and the detection line are displayed, signifying a negative result, whereas only the color of the quality control line indicates a positive result.

## Results

### The process of detecting schistosomiasis in rabbits

In this study, we combined RAA isothermal amplification technology with CRISPR-Cas12a to develop a method for detecting the infection of rabbits with *Schistosoma japonicum* (Fig. 1A). Nucleic acid was firstly extracted from a rabbit blood sample and then amplified with RAA at 37 ℃.

**Fig 1.**
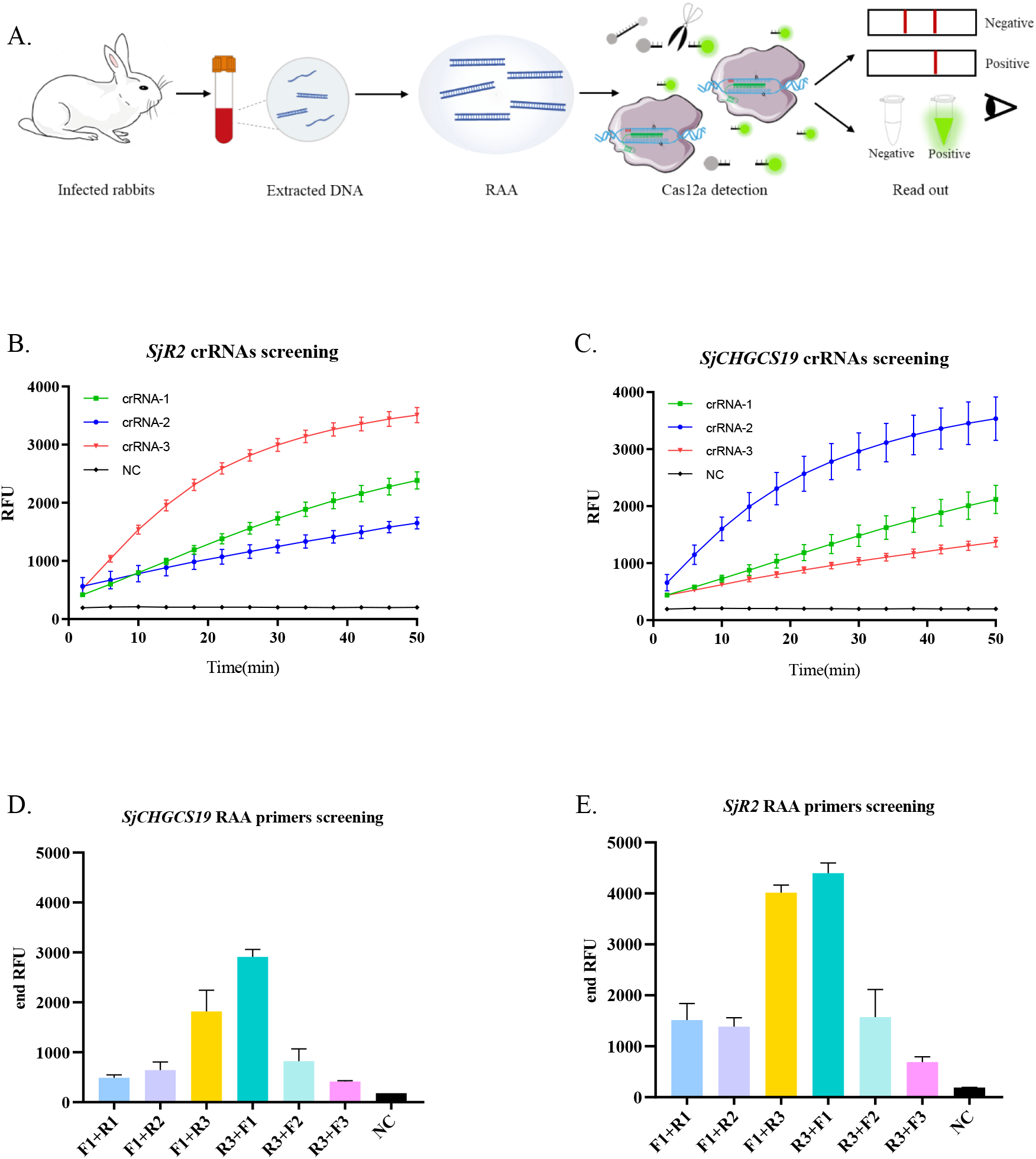
Screening of crRNA and RAA primers for CRISPR-Cas12a detection method. (A) The process of detecting schistosomiasis in rabbits. (B, C) The fluorescence signals of the *SjR2* and *SjCHGCS19* genes were detected during crRNA1, crRNA2, and crRNA3 screening. (D, E) Screening of RAA primers for *SjR2* and *SjCHGCS19* genes. (NC, RNase-free water)

The amplified DNA product were then added to the pre-prepared CRISPR-Cas12a reaction system, thereby activating the trans-cleavage ability of the Cas12a protein. The results can be detected through either fluorescence or lateral test strip.

### Design and screening of crRNA

This study aimed to detect circulating DNA in the serum of *Schistosoma japonicum*-infected rabbits. The two genes *SjR2* and *SjCHGCS19* of *Schistosoma japonicum* were chosen as target sites for the design of crRNA. Due to their multiple copies in the genome of *Schistosoma japonicum*, *SjR2* and *SjCHGCS19* are prone to generate more circulating DNA in serums for detection. Three crRNAs were designed for each of the conserved regions of *SjR2* and *SjCHGCS19*. The specificity of these crRNAs was confirmed using primer-blast in the NCBI database. To facilitate the screening of crRNA and RAA primers, plasmid standards of *SjR2* and *SjCHGCS19* were generated. We next assessed the performance of each crRNA to detect plasmid standards amplified with mixed RAA primers. The results showed that crRNA-3 in *SjR2* demonstrated the highest level of fluorescence. (Fig. 1B). The fluorescence signal of crRNA-2 was greatest in *SjCHGCS19* crRNA screening results (Fig. 1C).

### Screening of RAA primers

To improve the detection sensitivity of CRISPR-Cas12a, we performed a screening of RAA primers. Three sets of upstream and downstream primers were designed for *SjR2*-crRNA3 and *SjCHGCS19*-crRNA2, respectively. Next, several sets of RAA primers were screened using plasmid standards for two genes, *SjR2* and *SjCHGCS19*. Taking the gene *SjR2* as an example, first, use the forward primer to identify the best reverse primer by combining F1 with R1, R2, and R3, respectively, and in turn used the screened best reverse primer R3 to identify the most efficient forward primer. The results demonstrated that the primer combination of F1 + R3 had the best amplification effect (Fig. 1D). The same screening steps were performed on the RAA primers of the *SjCHGCS19* gene, and (Fig. 1E) showed that the F1 + R3 primer set exhibited the best amplification effect.

### CRISPR-Cas12a detection system optimization

In order to optimize the concentration of the reporter probe, we detected the effect of different concentrations of the DNA reporter probe on the fluorescence intensity of CRISPR-Cas12a protein detection. When the probe concentrations were: 160 nM, 80 nM, 40 nM, 20 nM, 10 nM, and 5 nM, the fluorescence intensity of the Cas12a detection result was positively correlated with the number of reporter probes within the detection range (Fig. 2A). Under the ultraviolet excitation, the fluorescence intensity of the solution in the tube increased proportionally with the concentration of the probe (Fig. 2B). These results indicate that increasing the number of reporter probes can improve the detection efficiency and signal intensity.

**Fig 2.**
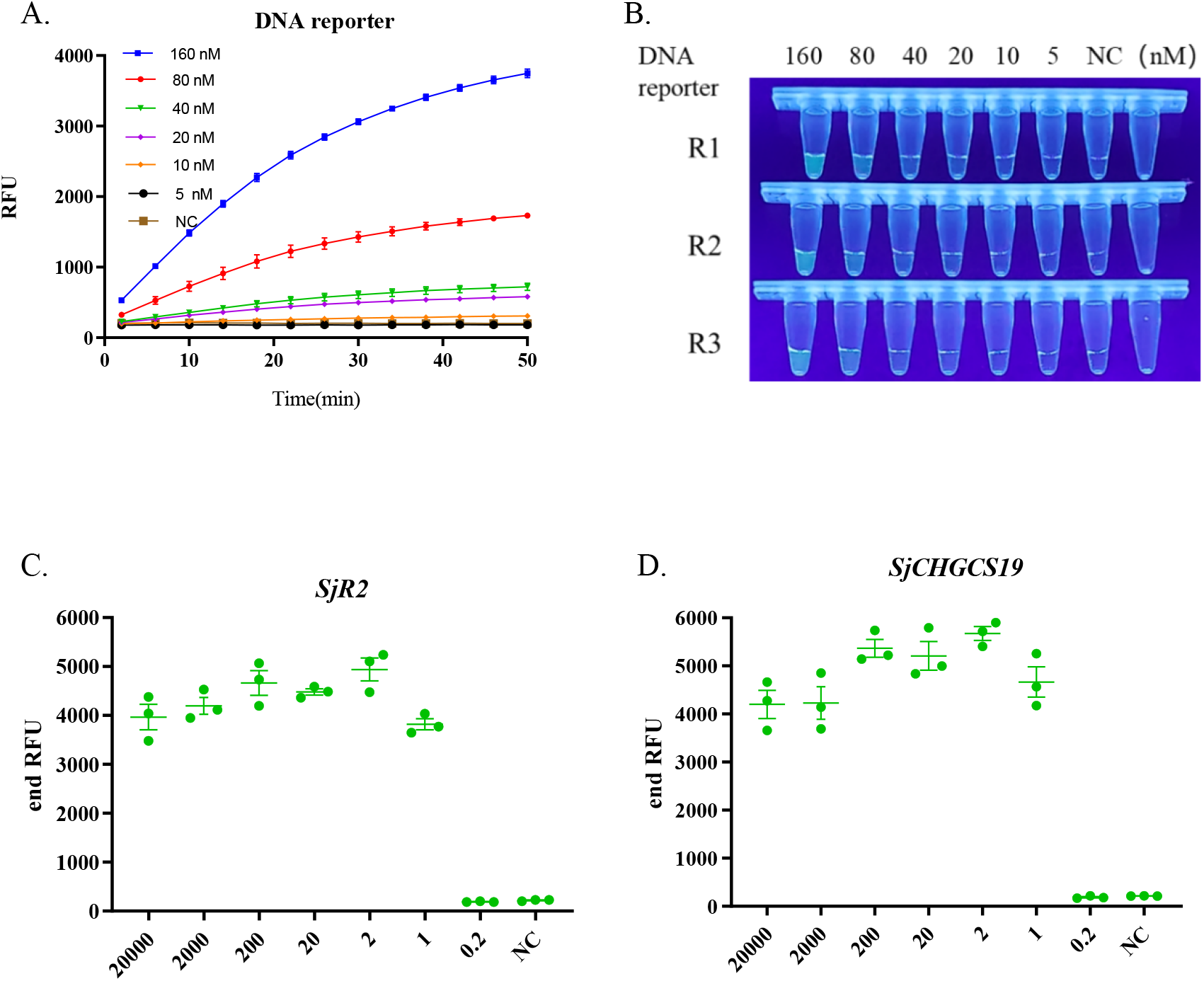
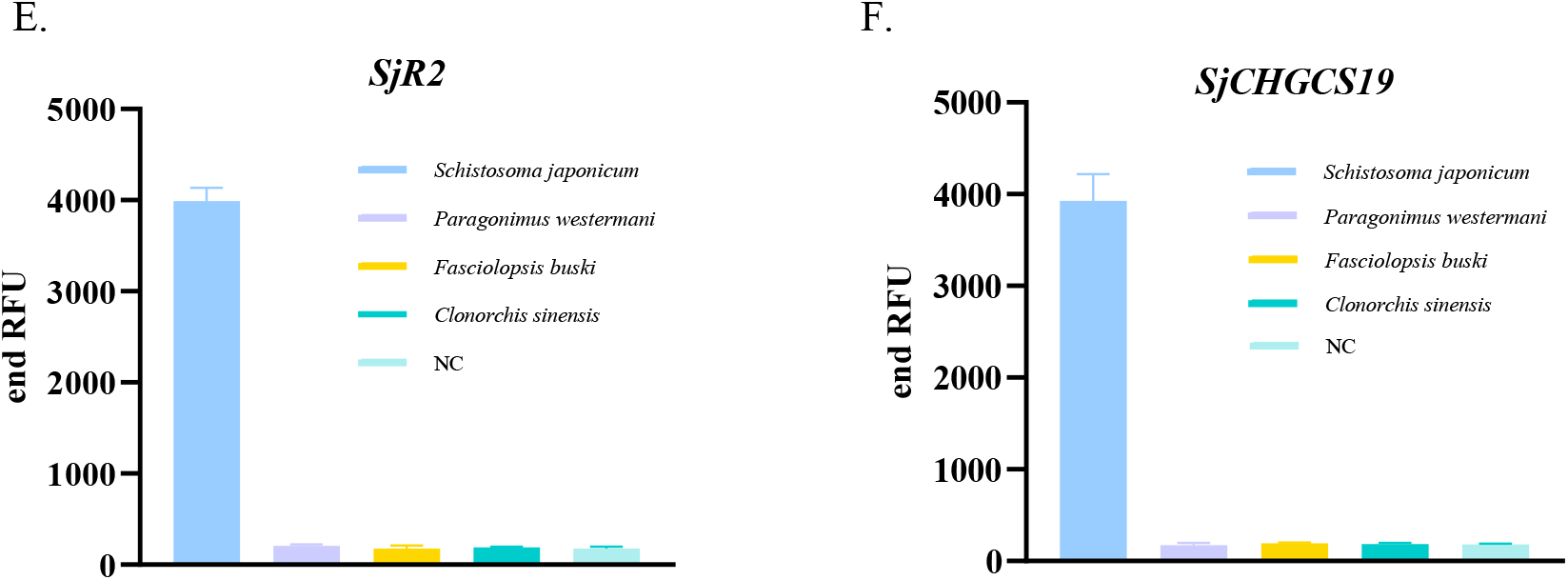
Optimization of CRISPR-Cas12a detection method and verification of sensitivity and specificity. (A, B) The system was optimized on DNA reporter concentration. CRISPR-Cas12a detected the real-time fluorescence signals of the plasmid standards when the probe concentration was 160 nM, 80 nM, 40 nM, 20 nM, 10 nM, and 5 nM, respectively. R1, R2, and R3 are abbreviations for three repetitions. (C, D) Gene detection sensitivity of CRISPR-Cas12a detection method for *SjR2* and *SjCHGCS19*. Serially diluted plasmid concentration was tested for each gene. 20000 cp/µL, 2000 cp/µL, 200 cp/µL, 20 cp/µL, 2 cp/µL, 1 cp/µL, and 0.2 cp/µL. (n=3, NC, RNase-free water.). (E, F) Specificity of the *SjR2* and *SjCHGCS19* CRISPR-Cas12a detection method. (n=3, NC, RNase-free water.)

### Sensitivity of CRISPR-Cas12a detection method

To verify the sensitivity of CRISPR-Cas12a for detection of the *SjR2* and *SjCHGCS19* genes, the constructed *SjR2* and *SjCHGCS19* gene plasmid standards were serially diluted into concentrations of 20000 cp/µL, 2000 cp/µL, 200 cp/µL, 20 cp/µL, 2 cp/µL, 1 cp/µL, and 0.2 cp/µL. The performance of each detection was assessed with the serially diluted plasmid standards. The results indicated that the detection sensitivity of SjR2 and SjCHGCS19 gene target sites was 1 cp/L. (Fig. 2C and D).

### Specificity of CRISPR-Cas12a detection method

To further assure the accuracy of the CRISPR-Cas12a method detection results, we validated the method’s specificity. First, the target sequences in the *SjR2* and *SjCHGCS19* genes were analyzed using primer-blast in the NCBI database, and sequence alignment revealed that the target sequences were specific. The CRISPR-Cas12a detection method was used to detect and analyze DNA samples from several other trematode parasites that infect humans, including *Schistosoma japonicum*, *Paragonimus westermani*, *Fasciolosis buski*, and *Clonorchis sinensis*. These results indicate that the CRISPR-Cas12a detection method was able to detect *Schistosoma japonicum* DNA without reacting with the DNA of other parasites (Fig. 2E and F).

### Comparison Nested-PCR and CRISPR-Cas12a detection methods for detecting the serum of rabbits infected with *Schistosoma japonicum*

To simulate the disease progression in patients with schistosomiasis, we made a rabbit infection model with 30 cercariae and collected serum at 1 d, 3 d, 1 w, 3 w, 5 w, and 7 w post-infection. Nested-PCR and CRISPR-Cas12a detection methods were applied to detect the samples. For Nested-PCR detection, 7 rabbits’ serums were detected positive for the *SjR2* gene (309 bp) on the third day after infection, with a positive rate of 47% (Fig. 3A); and 8 rabbits were detected positive for *the SjCHGCS19* gene (501 bp) with a positive rate of 53% (Fig. 3B) and can be detected continuously until the 7^th^ week.

**Fig 3.**
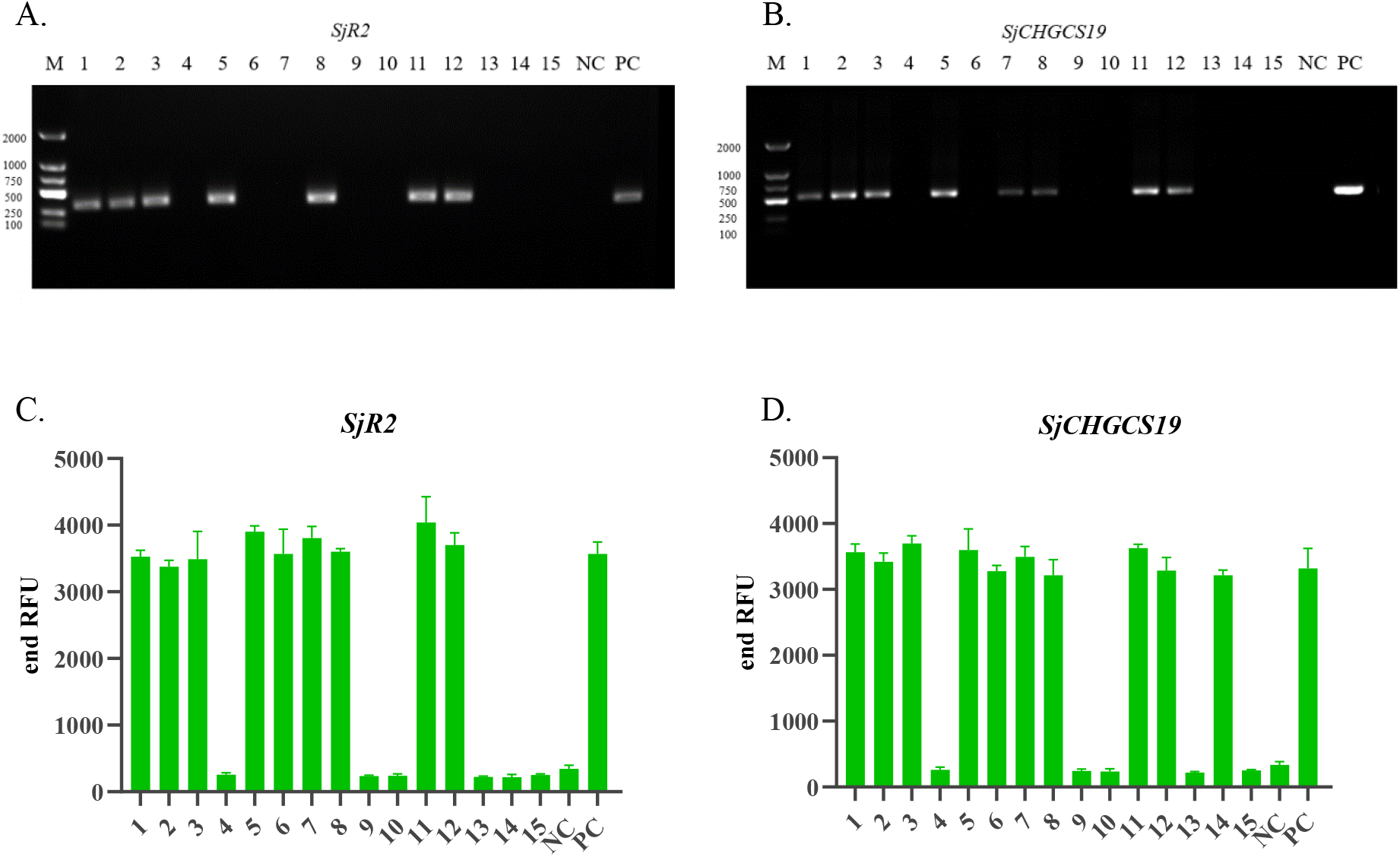
Nested-PCR and CRSIPR-Cas12a detection methods were used to detect the serum of rabbits infected with *Schistosoma japonicum*. (A, B) Electropherograms of *SjR2* and *SjCHGCS19* genes detected by Nested-PCR. (C, D) The CRSIPR-Cas12a method detected the relative fluorescence signals of *SjR2* and *SjCHGCS19* genes. (A, B, C, and D: 1-15 are serums from 15 rabbits infected with *Schistosoma japonicum*; NC: serums from uninfected rabbits; PC: *Schistosoma japonicum* genome).

For the CRISPR-Cas12a detection method, 9 rabbits were detected positive for the *SjR2* gene, with a positive rate of 60% (Fig. 3C); and 10 rabbits were detected positive for *SjCHGCS19* gene with a positive rate of 67% (Fig. 3D). The overall detection results were concluded in Table 1. Neither Nested-PCR nor the CRISPR-Cas12a detection method could detect the samples on 1^st^ d and 2^nd^ d post-infection, and the CRISPR-Cas12a detection method had higher positive rates than Nested-PCR on 3^rd^ d. Both methods could detect all infected samples at 3 weeks post-infection with a 100% positive rate.

**Table 1.**
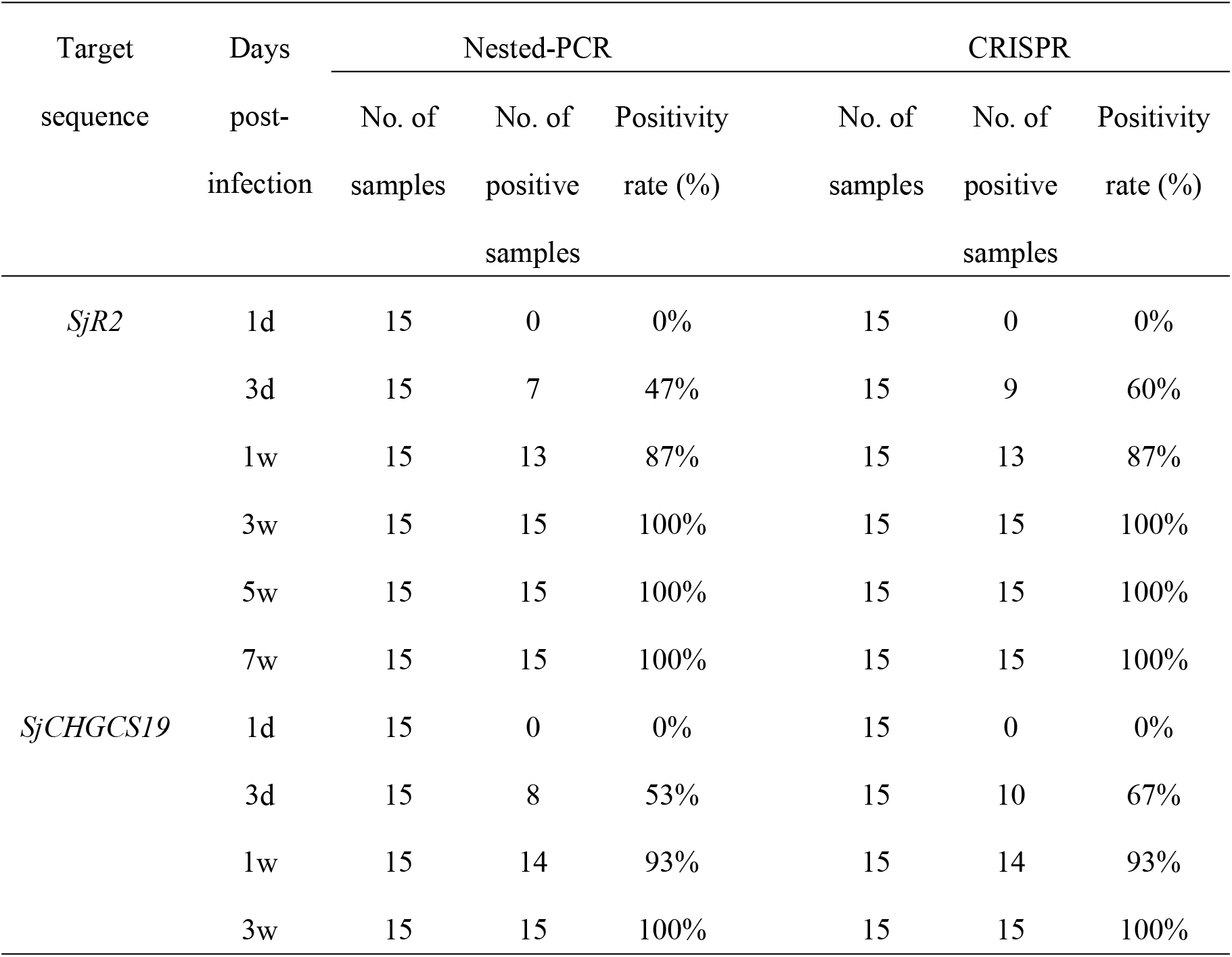
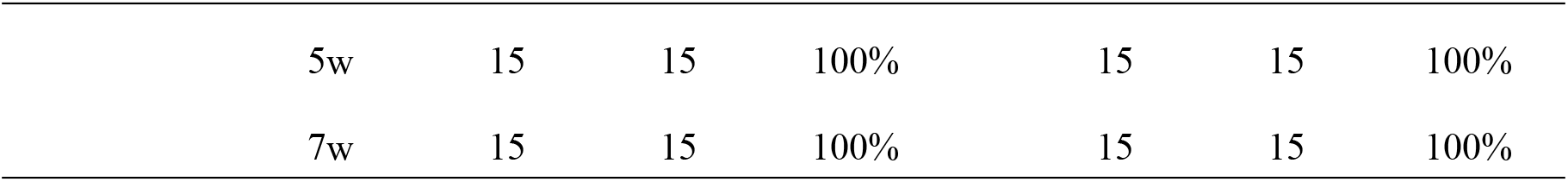
The positive rates of genes *SjR2* and *SjCHGCS19* detected by Nested-PCR and CRISPR-Cas12a detection methods at different time points of rabbit infection were compared.

### Field application of CRISPR-Cas12a detection method for detection of schistosomiasis

To make the detection method more accessible in remote areas with underdeveloped economies, this study utilized lateral test strips as the means of reading the results. In this study, the test strips will display two lines to indicate a negative result and one line for a positive result. The underlying principle is that the test strip sample binding pad contains a significant quantity of colloidal gold, which has FITC monoclonal antibody bound to it. (1) When there is no target DNA in the sample to be tested, the reporter probe (biotin-FAM) will form a complex with the colloidal gold bound to the FITC monoclonal antibody, and flow forward along the test strip. The biotin on the complex will be captured by streptomycin at the detection line, and the colloidal gold combined with biotin will develop color at the detection line. The remaining colloidal gold bound to FITC monoclonal antibody will continue to flow forward and be captured at the quality control line containing the anti-FITC antibody, and the quality control line will develop color. (2) When the sample to be tested contains the target DNA, CRSIPR-Cas12a protein will activate the non-specific trans-cleavage effect and cut the reporter probe (biotin-FAM) after recognizing the target DNA. Biotin without FAM cannot combine with colloidal gold, and the forward flow is captured by the detection line containing streptavidin, which does not develop color because it does not contain colloidal gold. The FAM-FITC antibody-colloidal gold complex was captured at the quality control line containing the anti-FITC antibody, and the quality control line developed color. The entire detection process can be completed within an hour and a half.

In this study, a test strip was used to confirm the specificity of the CRISPR-Cas12a detection method. The test strip can detect the *SjR2* and *SjCHGCS19* genes of *Schistosoma japonicum* without any cross-reactivity with other parasites, including *Paragonimus westermani*, *Fasciola brucei*, and *Clonorchis sinensis* (Fig. 4B and C).

**Fig 4.**
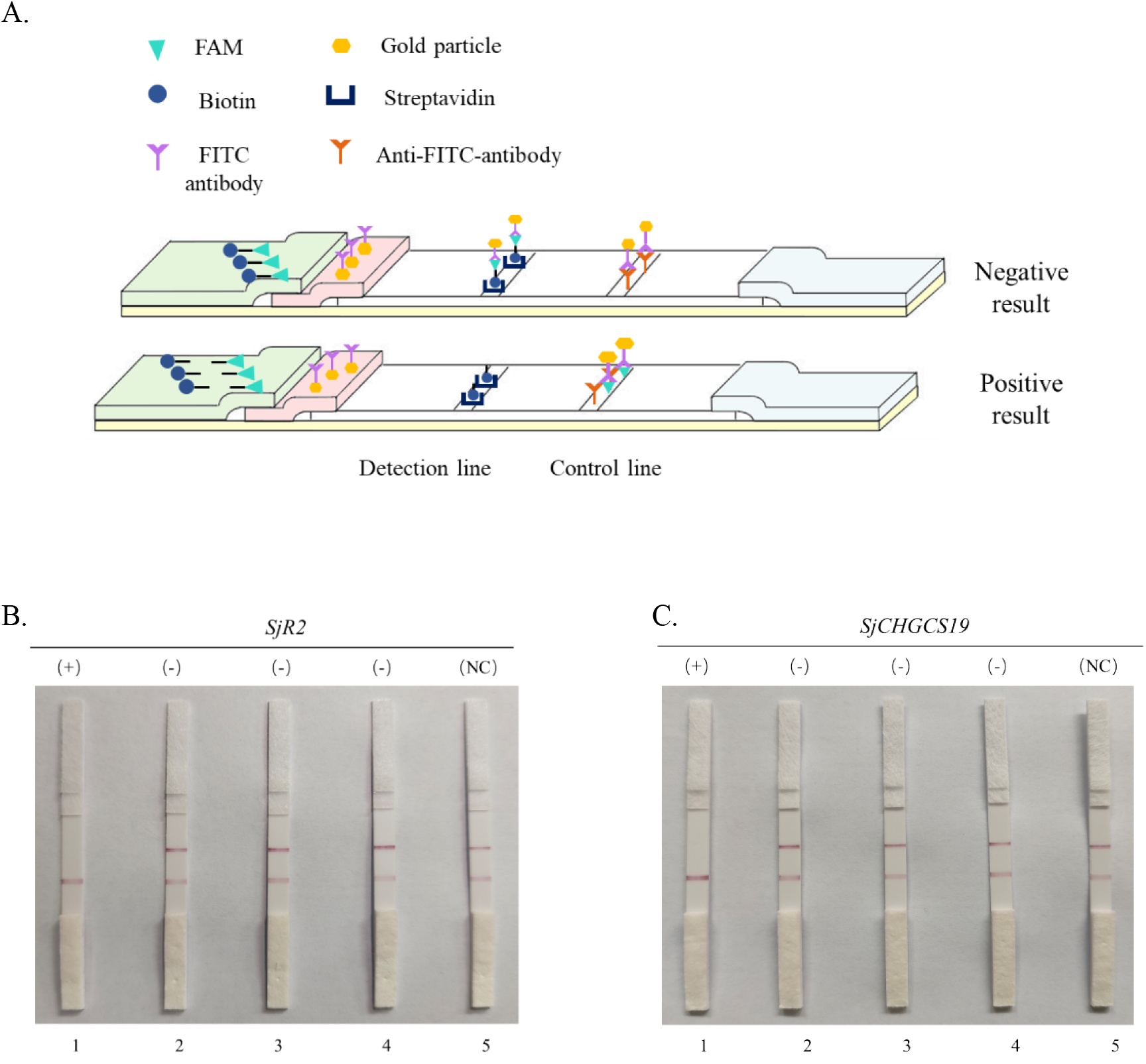
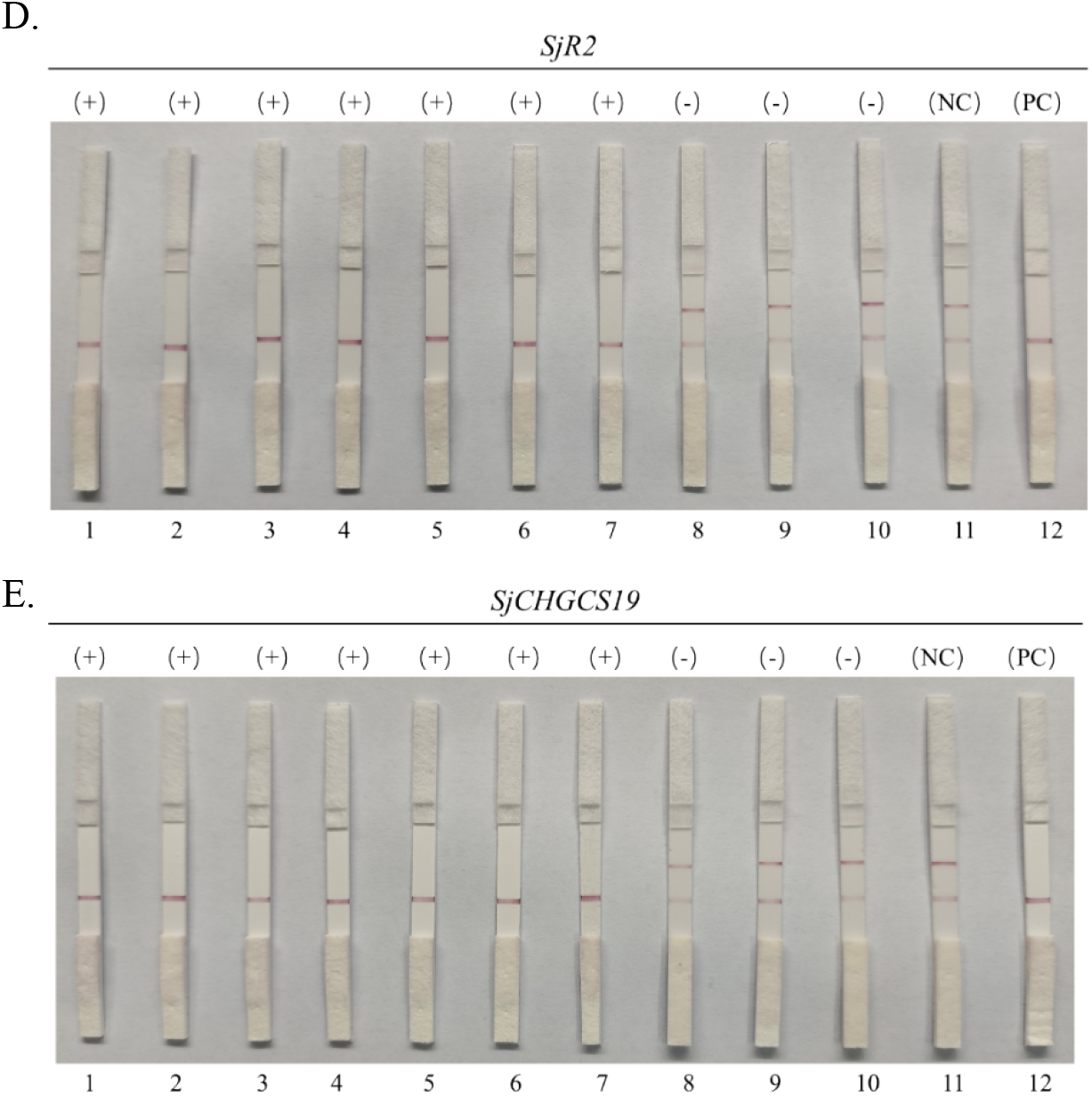
CRISPR-Cas12a detection method combined with lateral test strips for detection of *Schistosoma japonicum*. (A) Diagrammatic representation of the test strip detection principle. (B, C) Specificity of lateral test strips for detection of *SjR2* and *SjCHGCS19* genes in *Schistosoma japonicum*. (1-4 represent the *Schistosoma japonicum* DNA, *Paragonimus westermani* DNA, *Fasciolosis buski* DNA, and *Clonorchis sinensis* DNA, and 5 is a negative control adding RNase-free water. (D, E) Lateral test strips detect the *SjR2* and *SjCHGCS19* genes. (1-7 represent the serum from rabbits 3 weeks post-infection, 8-10 represent serum from uninfected rabbits, and 11 is a negative control for RNase-free water. 12 is a positive control within the genome of *Schistosoma japonicum*.)

To test the sensitivity of the test strip, serums from infected and healthy rabbits were used for detection. We chose 7 infected serums from rabbits 3 weeks post-infection and 3 un-infected serums from healthy rabbits to assess their reliability. The test strips were able to detect all 7 infected rabbits (Fig. 4D and E).

## Discussion

Schistosomiasis is listed by the World Health Organization as one of the 20 neglected tropical diseases and remains a significant public health concern globally[2,26]. The development of improved diagnostic methods for schistosomiasis is crucial to the control of this disease[27]. Considering the current focus on schistosomiasis control, it is necessary to establish a highly specific and sensitive diagnostic method that is also inexpensive and user-friendly. In this study, the RAA technique was selected as the signal amplification step for the nucleic acids under the examination. Because the temperature required for RAA is 37-45 ℃[28], which is consistent with the optimal reaction temperature of the CRISPR-Cas12a detection method, and the technique does not require complex equipments, it can quickly and efficiently complete isothermal amplification of the target within 20 minutes[29]. In this study, the RAA technique was combined with the CRISPR-Cas12a to establish a circulating nucleic acid detection technique for *Schistosoma japonicum*, and the experimental results were analyzed from multiple perspectives. In addition, the reaction conditions were optimized, and it was concluded that the sensitivity could reach the optimal reaction system for a single copy (1 cp/μL). In a rabbit model that simulates actual *Schistosoma japonicum* infection, the CRISPR-Cas12a detection method can detect the circulating nucleic acid of *Schistosoma japonicum* on the early 3rd day of infection, without cross-reactivity with other parasites. In addition, Cas12a was combined with a test strip in this study to make point-of-care testing (POCT). The entire detection process can be completed in one and a half hours at 37 ℃ or at room temperature without the use of sophisticated instruments. The CRISPR-Cas12a detection method is faster and more convenient than traditional PCR detection, meets the detection requirements of *Schistosoma japonicum* in the field environment, and enables immediate on-site diagnosis of schistosomiasis.

This study’s methodology consists of two parts: RAA amplification and Cas12a detection. The actual operation procedure must divide the amplification area from the sample input area to prevent aerosol contamination. Aman et al.[30]developed and optimized iSCAN-V2, a reverse transcription-recombinase polymerase amplification (RT-RPA) coupled CRISPR/Cas12b method for the detection of novel coronaviruses in a single reaction tube in less than one hour at a single temperature. Ding et al.[31] proposed a highly sensitive and specific dual CRISPR-Cas12a method called AIOD-CRISPR for detecting SARS-CoV-2. This method is straightforward and rapid, and it introduces dual crRNAs that are not limited by the original region adjacent motif (PAM) sequence, allowing for efficient dual CRISPR-based nucleic acid detection. The AIOD-CRISPR detection method is to add the components required for nucleic acid amplification and CRISPR detection into the same reaction tube, mix thoroughly, and incubate at constant temperature (37 ℃), which eliminates the separation of amplification products and transfer.

## Conclusion

In this study, specific target sites on the *Schistosoma japonicum* genes were identified, crRNA and RAA primers were designed. In conjunction with RAA isothermal amplification and the CRISPR-Cas12a detection method, a reliable detection method for schistosomiasis was established and the feasibility of the study was evaluated. The experimental results demonstrated that the method of this study has high sensitivity and specificity of a single copy, achieves low-cost, high-efficiency, real-time detection with naked-eye visualization, and solves the problem of equipment limitations, making it more appropriate for remote areas. Schistosomiasis detection on-site provides a new technical solution to the pressing problem of schistosomiasis detection. In addition, by designing various target sites for the detection of other pathogens, this method can provide new concepts for the detection of other pathogens.

## Acknowledgments

We express our gratitude to the Anhui Provincial Laboratory of Microbiology and Parasitology for their valuable support in providing the experimental platform. Additionally, we would like to extend our sincere appreciation to the Experimental Animal Research Center of the School of Basic Medicine, Anhui Medical University for their invaluable support.

## Author contributions

Conceptualization: Qian Li, Shuqi Yang, Jijia Shen, Yinan Du

Data curation: Qian Li, Shuqi Yang, Miao Cheng, Chen Xing

Formal analysis: Qian Li, Shuqi Yang, Miao Cheng

Funding acquisition: Yinan Du

Study: Qian Li, Shuqi Yang, Miao Cheng, Chen Xing, Xiaofeng Wang, Yating Zhu

Methodology: Qian Li, Shuqi Yang, Miao Cheng, Chen Xing, Yunshu Tang, Wenhui Cheng

Supervision: Jijia Shen, Yinan Du

Writing – original draft: Qian Li, Shuqi Yang

Writing – review & editing: Qian Li, Shuqi Yang, Miao Cheng, Chen Xing, Jijia Shen, Yinan Du

## Ethics statement

This study has been approved by the Experimental Animal Ethics Committee of Anhui Medical University (LLSC20221092, Anhui Medical University).

## Supporting information

**S1 Table.** Nested-PCR primers for the construction of standards.

**S2 Table.** RAA amplification primers.

**S3 Table.** DNA template sequences required for in vitro transcription of crRNA.

## References

1. Gryseels B, Polman K, Clerinx J, Kestens L. Human schistosomiasis.Lancet. 2006;368(9541):1106–18. Epub 2006/09/26. https://doi.org/10.1016/S0140-6736(06)69440-3 PubMed PMID: 16997665

2. McManus DP, Dunne DW, Sacko M, Utzinger J, Vennervald BJ, Zhou XN. Schistosomiasis. Nat Rev Dis Primers. 2018;4(1):13. Epub 2018/08/11. https://doi.org/10.1038/s41572-018-0013-8 PubMed PMID: 30093684

3. Weerakoon KG, Gobert GN, Cai P, McManus DP. Advances in the Diagnosis of Human S c-histosomiasis.Clin Microbiol Rev. 2015;28(4):939–67. Epub 2015/08/01. https://doi.org/10.1128/cmr.00137-14 PubMed PMID: 26224883

4. Molehin AJ. Schistosomiasis vaccine development: update on human clinical trials. J Biom-ed Sci. 2020;27(1):28. Epub 2020/01/24. https://doi.org/10.1186/s12929-020-0621-y.PubMed PMID: 31969170

5. Elmorshedy H, Bergquist R, Fayed A, Guirguis W, Abdel-Gawwad E, Eissa S, et al. Elimination of schistosomiasis requires multifactorial diagnostics: evidence from high- and low-prev-alence areas in the Nile Delta, Egypt. Infect Dis Poverty. 2020;9(1):31. https://doi.org/10.1186/s40249-020-00648-9 PubMed PMID: 32241298

6. Lo NC, Bezerra FSM, Colley DG, Fleming FM, Homeida M, Kabatereine N, et al. Revie-w of 2022 WHO guidelines on the control and elimination of schistosomiasis. Lancet Infect Dis. 2022;22(11):e327-e35. https://doi.org/10.1016/s1473-3099(22)00221-3. PubMed PMID: 35594896

7. Cavalcanti MG, Cunha AFA, Peralta JM. The Advances in Molecular and New Point-of-C-are (POC) Diagnosis of Schistosomiasis Pre- and Post-praziquantel Use: In the Pursuit of More Reliable Approaches for Low Endemic and Non-endemic Areas. Front Immunol. 2019;10:858. Epub 2019/06/14. https://doi.org/10.3389/fimmu.2019.00858. PubMed PMID: 31191512

8. Yu JM, de Vlas SJ, Jiang QW, Gryseels B. Comparison of the Kato-Katz technique, hatch-ing test and indirect hemagglutination assay (IHA) for the diagnosis of Schistosoma japonicum infection in China. Parasitol Int. 2007;56(1):45–9. https://doi.org/10.1016/j.parint.2006.11.002 Pu-bMed PMID: 17188018

9. Sarhan RM, Aminou HA, Saad GAR, Ahmed OA. Comparative analysis of the diagnostic performance of adult, cercarial and egg antigens assessed by ELISA, in the diagnosis of chronic human Schistosoma mansoni infection. Parasitol Res. 2014;113(9):3467–76. https://doi.org/10.1007/s00436-014-4017-3. PubMed PMID: 25028207

10. Bärenbold O, Garba A, Colley DG, Fleming FM, Assaré RK, Tukahebwa EM, et al. Estimating true prevalence of Schistosoma mansoni from population summary measures based on the Kato-Katz diagnostic technique. PLoS Negl Trop Dis. 2021;15(4):e0009310. https://doi.org/10.1371/journal.pntd.0009310. PubMed PMID: 33819266

11. Yu JM, de Vlas SJ, Yuan HC, Gryseels B. Variations in fecal Schistosoma japonicum egg counts. Am J Trop Med Hyg. 1998;59(3):370–5. https://doi.org/10.4269/ajtmh.1998.59.370 PubMed PMID: 9749627

12. Utzinger J, Becker SL, van Lieshout L, van Dam GJ, Knopp S. New diagnostic tools in s-chistosomiasis. Clin Microbiol Infect. 2015;21(6):529–42. https://doi.org/10.1016/j.cmi.2015.03.014. PubMed PMID: 25843503

13. Yamano T, Nishimasu H, Zetsche B, Hirano H, Slaymaker IM, Li Y, et al. Crystal Structure of Cpf1 in Complex with Guide RNA and Target DNA. Cell. 2016;165(4):949–62. https://doi.org/10.1016/j.cell.2016.04.003. PubMed PMID: 27114038

14. Zetsche B, Gootenberg JS, Abudayyeh OO, Slaymaker IM, Makarova KS, Essletzbichler P, et al. Cpf1 is a single RNA-guided endonuclease of a class 2 CRISPR-Cas system. Cell. 2015;163(3):759–71. https://doi.org/10.1016/j.cell.2015.09.038. PubMed PMID: 26422227

15. Deng H, Gao Z. Bioanalytical applications of isothermal nucleic acid amplification techniques. Anal Chim Acta. 2015;853:30–45. https://doi.org/10.1016/j.aca.2014.09.037. PubMed PMID: 25467448

16. Chen JS, Ma E, Harrington LB, Da Costa M, Tian X, Palefsky JM, et al. CRISPR-Cas12a target binding unleashes indiscriminate single-stranded DNase activity. Science. 2018;360(6387):436-9. https://doi.org/10.1126/science.aar6245. PubMed PMID: 29449511

17. Liu X, Qiu X, Xu S, Che Y, Han L, Kang Y, et al. A CRISPR-Cas12a-Assisted Fluorescence Platform for Rapid and Accurate Detection of Nocardia cyriacigeorgica. Front Cell Infect Microbiol. 2022;12:835213. Epub 2022/03/22. https://doi.org/10.3389/fcimb.2022.835213. PubM-ed PMID: 35310854

18. Xu H, Tang H, Li R, Xia Z, Yang W, Zhu Y, et al. A New Method Based on LAMP-CRISPR-Cas12a-Lateral Flow Immunochromatographic Strip for Detection. Infect Drug Resist. 2022;15:685–96. Epub 2022/03/08. https://doi.org/10.2147/idr.s348456. PubMed PMID: 35250283

19. Xing S, Lu Z, Huang Q, Li H, Wang Y, Lai Y, et al. An ultrasensitive hybridization chain reaction-amplified CRISPR-Cas12a aptasensor for extracellular vesicle surface protein quantification. Theranostics. 2020;10(22):10262–73. Epub 2020/09/16. https://doi.org/10.7150/thno.49047. PubMed PMID: 32929347

20. Ma QN, Wang M, Zheng LB, Lin ZQ, Ehsan M, Xiao XX, et al. RAA-Cas12a-Tg: A Nucleic Acid Detection System for Toxoplasma gondii Based on CRISPR-Cas12a Combined withRecombinase Aided Amplification (RAA). Microorganisms. 2021;9(8). Epub 2021/08/28. https://doi.org/10.3390/microorganisms9081644. PubMed PMID: 34442722

21. Lei R, Li L, Wu P, Fei X, Zhang Y, Wang J, et al. RPA/CRISPR/Cas12a-Based On-Site and Rapid Nucleic Acid Detection of Toxoplasma gondii in the Environment. ACS Synth Biol. 2022;11(5):1772–81. Epub 2022/04/27. https://doi.org/10.1021/acssynbio.1c00620.1c00620. PubMe-d PMID: 35471824

22. Shen X-X, Qiu F-Z, Shen L-P, Yan T-F, Zhao M-C, Qi J-J, et al. A rapid and sensitive recombinase aided amplification assay to detect hepatitis B virus without DNA extraction. BM-C Infect Dis. 2019;19(1):229. https://doi.org/10.1186/s12879-019-3814-9. PubMed PMID: 30836947

23. Laha T, Brindley PJ, Smout MJ, Verity CK, McManus DP, Loukas A. Reverse transcripta s-e activity and untranslated region sharing of a new RTE-like, non-long terminal repeat r et-rotransposon from the human blood fluke, Schistosoma japonicum. Int J Parasitol. 2002; 32(9):1163–74. https://doi.org/10.1016/s0020-7519(02)00063-2 PubMed PMID: 12117499

24. Guo JJ, Zheng HJ, Xu J, Zhu XQ, Wang SY, Xia CM. Sensitive and specific target sequences selected from retrotransposons of Schistosoma japonicum for the diagnosis of schist o-somiasis. PLoS Negl Trop Dis. 2012;6(3):e1579. Epub 2012/04/06. https://doi.org/10.1371/journal.pntd.0001579. PubMed PMID: 22479661

25. Ullah H, Qadeer A, Giri BR. Detection of circulating cell-free DNA to diagnose Schistosoma japonicum infection. Acta Trop. 2020;211:105604. Epub 2020/07/01. https://doi.org/10.1016/j.actatropica.2020.105604. PubMed PMID: 32598919

26. Meltzer E. Schistosomiasis: still a neglected disease. J Travel Med. 2021;28(6). https://doi.org/10.1093/jtm/taab107. PubMed PMID: 34254141

27. Cavalcanti MG, Silva LF, Peralta RHS, Barreto MGM, Peralta JM. Schistosomiasis in areas of low endemicity: a new era in diagnosis. Trends In Parasitology. 2013;29(2):75–82. https://doi.org/10.1016/j.pt.2012.11.003. PubMed PMID: 23290589

28. Xing W, Yu X, Feng J, Sun K, Fu W, Wang Y, et al. Field evaluation of a recombinase polymerase amplification assay for the diagnosis of Schistosoma japonicum infection in Hunan province of China. BMC Infect Dis. 2017;17(1):164. https://doi.org/10.1186/s12879-017-2182-6. PubMed PMID: 28222680

29. Lobato IM, O’Sullivan CK. Recombinase polymerase amplification: Basics, applications and recent advances. Trends Analyt Chem. 2018;98:19–35. https://doi.org/10.1016/j.trac.2017.10.015. PubMed PMID: 32287544

30. Aman R, Marsic T, Sivakrishna Rao G, Mahas A, Ali Z, Alsanea M, et al. iSCAN-V2: A One-Pot RT-RPA-CRISPR/Cas12b Assay for Point-of-Care SARS-CoV-2 Detection. Front Bio-eng Biotechnol. 2021;9:800104. Epub 2022/02/08. https://doi.org/10.3389/fbioe.2021.800104. Pub-Med PMID: 35127671

31. Ding X, Yin K, Li Z, Lalla RV, Ballesteros E, Sfeir MM, et al. Ultrasensitive and visual detection of SARS-CoV-2 using all-in-one dual CRISPR-Cas12a assay. Nat Commun. 2020;11(1):4711. Epub 2020/09/20. https://doi.org/10.1038/s41467-020-18575-6. PubMed PMID: 32948757

